# Hypoxia inducible factors regulate infectious SARS-CoV-2, epithelial damage and respiratory symptoms in a hamster COVID-19 model

**DOI:** 10.1101/2022.03.15.484379

**Authors:** Peter A.C. Wing, Maria Prange-Barczynska, Amy Cross, Stefania Crotta, Claudia Orbegozo Rubio, Xiaotong Cheng, James M. Harris, Xiaodong Zhuang, Rachel L. Johnson, Kathryn A. Ryan, Yper Hall, Miles W. Carroll, Fadi Issa, Peter Balfe, Andreas Wack, Tammie Bishop, Francisco J. Salguero, Jane A. McKeating

**Author notes:** Shared Senior Authorship (Francisco J. Salguero, Jane A. McKeating). Corresponding authors, Corresponding authors: Jane A McKeating and Peter AC Wing.

## Abstract

Understanding the host pathways that define susceptibility to SARS-CoV-2 infection and disease are essential for the design of new therapies. Oxygen levels in the microenvironment define the transcriptional landscape, however the influence of hypoxia on virus replication and disease in animal models is not well understood. In this study, we identify a role for the hypoxic inducible factor (HIF) signalling axis to inhibit SARS-CoV-2 infection, epithelial damage and respiratory symptoms in Syrian hamsters. Pharmacological activation of HIF with the prolyl-hydroxylase inhibitor FG-4592 significantly reduced the levels of infectious virus in the upper and lower respiratory tract. Nasal and lung epithelia showed a reduction in SARS-CoV-2 RNA and nucleocapsid expression in treated animals. Transcriptomic and pathological analysis showed reduced epithelial damage and increased expression of ciliated cells. Our study provides new insights on the intrinsic antiviral properties of the HIF signalling pathway in SARS-CoV-2 replication that may be applicable to other respiratory pathogens and identifies new therapeutic opportunities.

## INTRODUCTION

COVID-19, caused by the coronavirus SARS-CoV-2, is a global health issue with more than 5.5 million fatalities to date. Vaccination has reduced both the number of hospitalisations and mortality due to COVID-19 (Singanayagam et al., 2021; Voysey et al., 2021). However, the emergence of variants, such as Omicron, that show reduced sensitivity to vaccine-induced immunity (Dejnirattisai et al., 2021a; Dejnirattisai et al., 2021b; Liu et al., 2021), provide the potential for new waves of infection. The primary site of SARS-CoV-2 infection is the upper respiratory epithelia with diminishing levels of infection in distal areas of the lung (Hou et al., 2020). A defining feature of severe COVID-19 pneumonitis is systemic low oxygen (hypoxaemia), which can lead to organ failure and death through acute respiratory distress syndrome (Huang et al., 2020; Li et al., 2020). At the cellular level, hypoxia induces substantial changes to the host transcriptional landscape regulating a diverse array of biological pathways that are orchestrated by hypoxic inducible factors (HIFs). When oxygen is abundant, newly synthesised HIFα subunits are hydroxylated by HIF prolyl-hydroxylase domain (PHD) enzymes resulting in their proteasomal degradation. Under hypoxic conditions the PHD enzymes are inactive and stabilised HIFα dimerizes with HIF-1β, translocates to the nucleus, and promotes the transcription of genes involved in erythropoiesis, glycolysis, pulmonary vasomotor control, and immune regulation (Kaelin and Ratcliffe, 2008; Palazon et al., 2014; Urrutia and Aragones, 2018). HIF-target genes can vary between cell types allowing a flexible response to diverse physiological signals (Schodel et al., 2011).

Under normal physiological conditions, the lungs provide an oxygen rich environment, however, an increasing body of literature shows a role for hypoxia in the inflamed airway epithelium (Page et al., 2021). Transcriptomic analysis of post-mortem COVID-19 pulmonary tissue shows an association between hypoxic signalling and inflammatory responses (Cross et al., 2021; Sposito et al., 2021). While HIFs may drive inflammation in certain settings, HIF-1α has been shown to suppress the inflammatory response in bronchial epithelial cells reducing expression of IL-6 and IP10 (Polke et al., 2017). This dual role of HIFs highlights the importance of the cellular environment in which hypoxia occurs.

HIFs modulate the replication of a wide number of viruses (Liu et al., 2020), enhancing the replication of hepatitis B (Wing et al., 2021b) and Epstein Barr viruses (Jiang et al., 2006; Kraus et al., 2017) via direct binding to their viral DNA genomes. In contrast, HIFs inhibit influenza A virus replication in lung epithelial models of infection (Zhao et al., 2020). These differing outcomes may reflect variable oxygen levels at the site of virus replication in the body. Several respiratory pathogens including Influenza (Ren et al., 2019), Rhinovirus (Gualdoni et al., 2018) and Respiratory Syncytial virus (Haeberle et al., 2008) induce anaerobic glycolysis via activation of the HIF-1α signalling axis, suggesting a role for viruses to manipulate this pathway. A greater understanding of the oxygen microenvironment in the healthy and inflamed lung will inform our understanding of mucosal host-pathogen interactions.

We have reported that hypoxic activation of HIF-1α inhibits SARS-CoV-2 entry and replication in primary and immortalised lung epithelial cells (Wing et al., 2021a). HIF-1α downregulates the expression of two key entry factors ACE2 and TMPRSS2, thereby limiting SARS-CoV-2 internalisation, whilst also restricting the establishment of viral replication complexes. These data show an essential role for hypoxia/HIF-1α in multiple aspects of the SARS-CoV-2 life cycle and it is timely to address the role of HIFs in an immune competent animal model of COVID-19 disease.

HIFs can be activated by drugs that inhibit the PHDs which are currently used for the treatment of renal anaemia (Akizawa et al., 2020a; Akizawa et al., 2020b; Akizawa et al., 2020c; Akizawa et al., 2020d; Chen et al., 2019a; Chen et al., 2019b). We evaluated the ability of the PHD inhibitor FG-4592 (Roxadustat) to inhibit SARS-CoV-2 replication and pathogenesis in Golden Syrian hamsters, that shows similar features to human disease including lung pathology and damage to the ciliated epithelia (Chan et al., 2020; de Melo et al., 2021; Imai et al., 2020; Rosenke et al., 2020; Sia et al., 2020). Treatment of infected hamsters with FG-4592, either prophylactically or after infection, reduced the infectious viral burden and respiratory symptoms. Our study provides new information on how the HIF signalling pathway influences SARS-CoV-2 replication that may be applicable to other respiratory pathogens and suggests new preventative and therapeutic opportunities

## RESULTS

### Orally administered FG-4592 activates HIFs in the lung and limits SARS-CoV-2 disease severity

To assess the effect of FG-4592 on SARS-CoV-2 infection, hamsters were treated with 30mg/kg of drug twice daily by oral gavage commencing either 24h pre- or 24h post-viral challenge. This regimen was based on previous FG-4592 dosing protocols in mice (Schley et al., 2019; Wing et al., 2021a) and clinical studies (Provenzano et al., 2016). In the control group, animals were treated with vehicle 24h prior to infection which continued throughout the study in the same manner as treated animals **(Fig.1A).** Hamsters were infected with SARS-CoV-2 (Australia/VIC01/2020 or VIC01) by intranasal delivery of 5×10^4^ plaque forming units (PFU), which is sufficient to cause clinical signs and respiratory lesions (Huo et al., 2021; Rosenke et al., 2020; Ryan et al., 2021). Weight and body temperature were recorded and clinical signs such as laboured breathing, ruffled fur and lethargy measured twice daily to provide a clinical score (described in **Supplementary Table 1**). Infectious virus in the upper respiratory tract was measured in nasal washes and throat swabs collected at 1-, 2- and 4-days post-infection. The study was terminated at 4 days post-infection based on studies reporting the detection of infectious virus in the upper respiratory tract **(Fig.1A)** (Chan et al., 2020; Imai et al., 2020; Rosenke et al., 2020).

**Figure 1:**
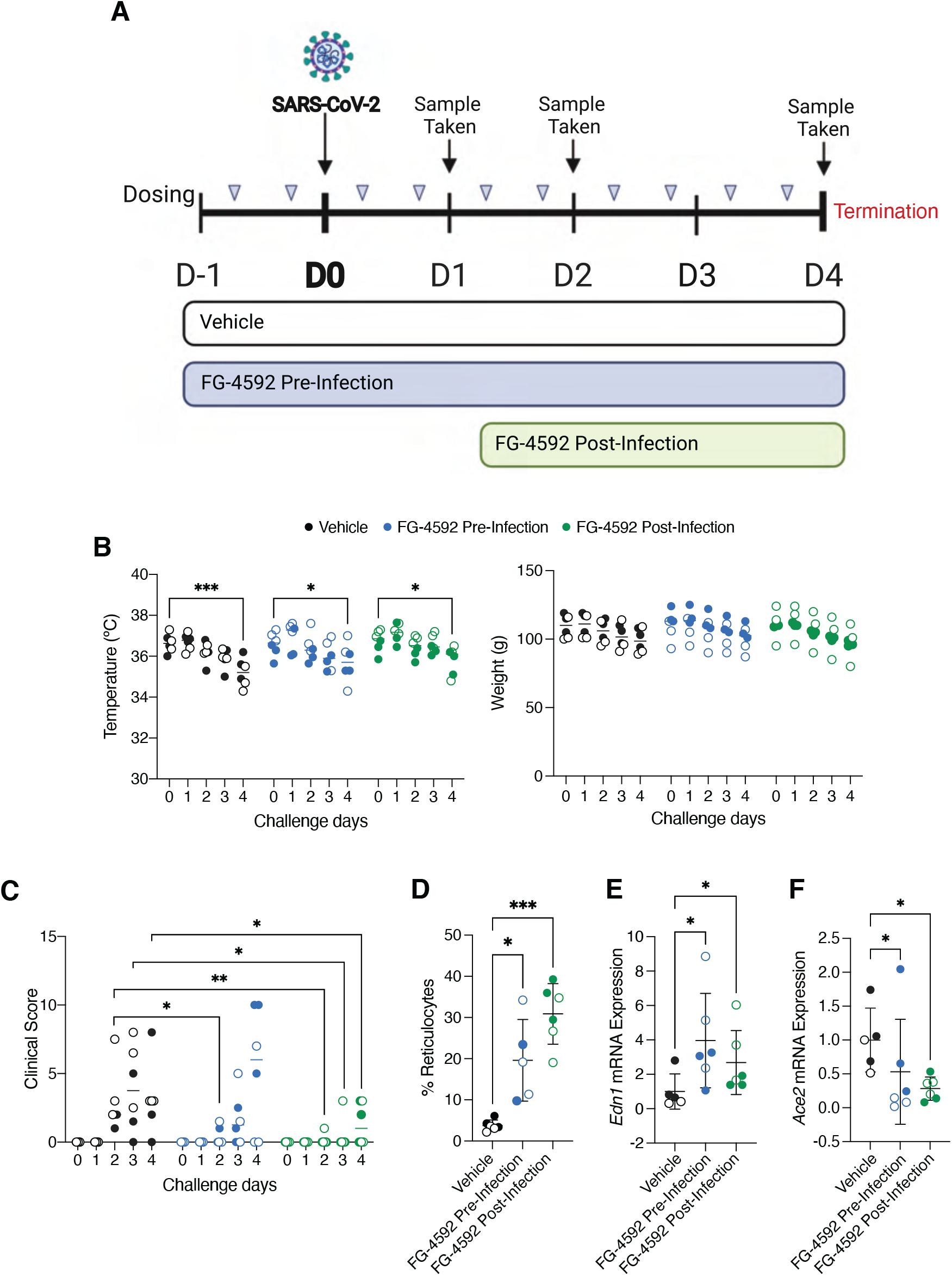
Evaluating FG-4592 in the Syrian hamster model of SARS-CoV-2. **(A)** Schematic of the challenge study. 18 animals aged 7 weeks were allocated into groups of 6 for each treatment. For vehicle and pre-infection groups, twice daily dosing of vehicle or 30mg/kg of FG-4592 administered by oral gavage commenced 24h prior to intranasal challenge with SARS-CoV-2 VIC01/2020 (5×10^4^ PFU). For the post-infection treatment group, FG-4592 dosing started 24h after viral infection. On days 1, 2 and 4 post-infection, nasal washes and throat swabs were collected for assessment of viral load and infectious titre. The study was terminated at 4 days post-infection. **(B)** Hamster temperature and weight measurements over the course of the study. Weights were measured daily, and temperature recorded twice daily with average measurements for each animal plotted. **(C)** Clinical scores from daily animal assessments across the treatment groups. Observations such as wasp waisted, ruffled fur, hunched or laboured breathing were recorded and assigned a numerical value (see **Supplementary Table 1**). **(D)** Reticulocyte counts were quantified by staining terminal blood samples with 0.1% Brilliant Cresyl Blue. Endothelin-1 (*Edn1*) **(E)** and *Ace2* **(F)** mRNA levels in the lung and data expressed relative to the mean of the control Vehicle group. Open circles represent female animals and closed circles males. Unless otherwise stated, data is expressed as mean ± s.d. Statistical analysis was performed using a one-way ANOVA, p<0.05 = *, p<0.01 = **, p<0.001 = ***.

We observed a significant reduction in body temperature and a loss of body weight in all treatment groups **(Fig.1B)**, in line with the clinical presentation of SARS-CoV-2 in this model (Chan et al., 2020; Nouailles et al., 2021; Rosenke et al., 2020). No significant differences in animal weight or temperature were noted between the treatment groups, suggesting that FG-4592 was well-tolerated **(Fig.1B).** The first signs of disease were observed at day 2 post-infection, further increasing by day 4 in the control group primarily due to the onset of laboured breathing **(Fig.1C, Supplementary Table 1)**. Animals treated with FG-4592 showed a significant improvement in their clinical score, particularly in the post-infection treatment group **(Fig.1C)**. Further, while all animals in the control vehicle group presented with laboured breathing, this was only observed in 2/6 animals in the pre-infection treatment group and none of the hamsters in the post-infection treatment group.

As HIF expression following systemic PHI treatment is transient and difficult to detect (Chan et al., 2016), we evaluated FG-4592 efficacy by assessing HIF activation of erythropoietin stimulated erythrocytosis by measuring immature red blood cells (reticulocytes). Blood smears from terminal blood samples showed increased reticulocyte counts compared to vehicle, consistent with effective drug treatment **(Fig.1D)**. To evaluate whether FG-4592 activated HIFs in the lung we assessed pulmonary expression of the HIF target gene Endothelin-1 (*Edn-1*) (Hickey et al., 2010) and noted a modest but significant induction of mRNA (**Fig.1E**). Furthermore, we noted a decrease in mRNA levels of the viral entry receptor *Ace2* in the lungs of treated hamsters **(Fig.1F),** supporting our previous findings (Wing et al., 2021a). To understand the PHI-driven changes in pulmonary gene expression we sequenced RNA from the lung tissue of vehicle, FG-4592 pre- or post-infection groups and observed an induction of 47 and 63 genes respectively, including HIF target genes such as *Edn-1* and *Bnip3* **(Supplementary Fig.1A)**. To assess whether all animals responded to FG-4592 we evaluated transcript levels of the common HIF-upregulated genes. Hierarchical cluster analysis separated the vehicle and treated animals and showed comparable activation in the pre- and post-infection treatment groups, demonstrating that animals had responded in a similar manner **(Supplementary Fig.1B)**. Together these data show that FG-4592 is well tolerated, activates HIF-transcriptional responses in the lung and reduces symptoms of SARS-CoV-2 infection.

### FG-4592 reduces infectious SARS-CoV-2 in upper and lower respiratory tract

The course of SARS-CoV-2 disease in the Syrian hamster is transient, with the onset of clinical symptoms peaking between 4-6 days post-infection followed by the development of neutralising antibodies and viral clearance within 8-15 days (Chan et al., 2020; Sia et al., 2020). To assess the effect of HIFs on SARS-CoV-2 replication we measured viral RNA by qPCR and infectious virus by plaque assay using Vero-TMPRSS2 cells **(Supplementary Fig.2)**. High levels of viral RNA and infectious SARS-CoV-2 were detected in the nasal washes and throat swabs sampled at day 1 post-infection in the vehicle group, which declined over the course of the study **(Fig.2A-B)**. Pre-treatment with FG-4592 resulted in a 1-log reduction in the infectious viral burden in both nasal washes and throat swabs at day 2 post-infection **(Fig.2A-B)**. Similarly, animals treated post-infection showed significantly reduced levels of infectious virus by day 4. In contrast, drug treatment had a negligible effect on the total viral RNA levels measured in either the nasal washes or throat swabs **(Fig.2A-B).** We also measured the burden of infectious virus in the lungs at the end of the study and showed a significant reduction in the treated animals (**Fig.2C).** However, there was no substantial change in total or genomic viral RNA (gRNA) **(Fig.2C, Supplementary Fig.3)**. Together these data demonstrate that PHI treatment before or after infection significantly reduced the infectious viral burden in the upper and lower respiratory tract of infected hamsters.

**Figure 2:**
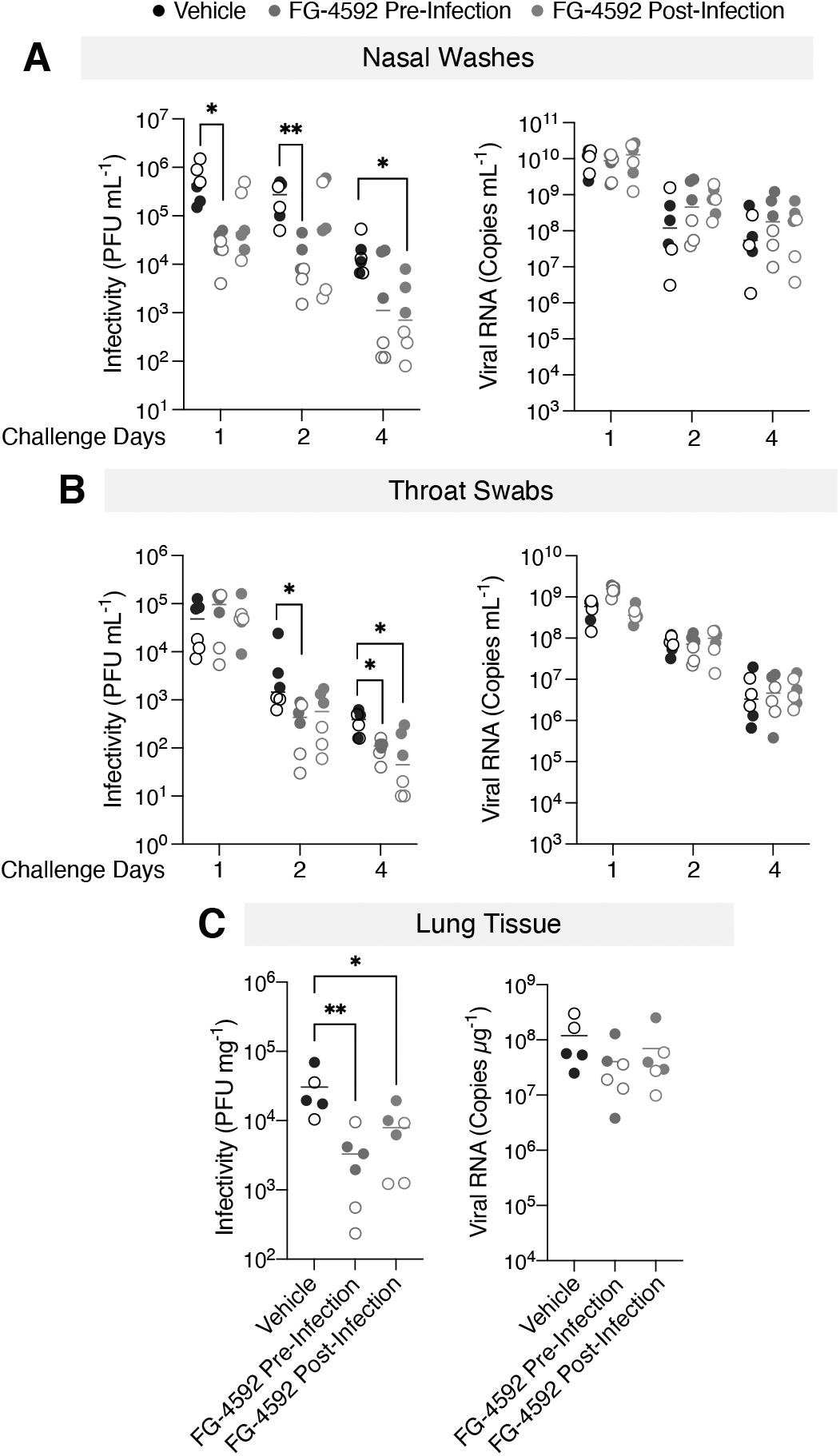
FG-4592 treatment reduces the infectious SARS-CoV-2 burden in the respiratory tract. **(A)** The infectious burden in the nasal washes sampled at days 1, 2, and 4 post-infection was quantified by plaque assay on Vero-TMPRSS2 cells and data presented as plaque forming units (PFU) per ml. Viral RNA copies were measured by RT-qPCR and expressed as copies/mL. **(B)** Viral infectivity and RNA copies were measured in throat swabs as described above. **(C)** Snap frozen lung samples from the right lobe harvested at termination were homogenised for RNA extraction and titration of infectious virus. Viral RNA copies were quantified by RT-qPCR and expressed as copies/μg of total RNA. Infectious titre was determined by plaque assay and expressed as PFU/mg of lung tissue. Open circles represent female animals and closed circles males. Statistical analysis was performed by ANOVA, p<0.05 = *, p<0.01 = **, p<0.0001 = ***, p<0.0001 = ****. Brackets indicate the comparisons tested.

### FG-4592 reduces SARS-CoV-2 sub-genomic RNAs in the lung

Since FG-4592 reduced the level of infectious virus in the lung we were interested to assess whether treatment impacts the viral transcriptome. Mapping the viral reads across the 30kb SARS-CoV-2 genome demonstrated an increasing read depth from ORF1ab to the 3’UTR consistent with the transcription of sub-genomic (sg) RNAs **(Supplementary Fig.4A).** In addition to the gRNA, the viral transcriptome includes 9 canonical sub-genomic (sg) RNAs that encode the structural proteins, which are essential for the genesis of nascent virus particles. Quantifying the junction spanning reads between the common 5’ leader sequence and the start of each sgRNA (as previously described (Kim et al., 2020)), enabled us to infer their approximate abundance. FG-4592 reduced the abundance of most sgRNAs with a greater variability in the treated groups and a significant reduction in the nucleocapsid (N) transcript, the most abundant of the viral RNAs **(Fig.3A)**. We extended these observations to study the effect of HIF-signalling in SARS-CoV-2 transcription in the lung Calu-3 epithelial cell line (Sampaio et al., 2021) and showed a significant reduction in S, E, M, ORF6, ORF7A, ORF7B, ORF8 and N junction spanning reads **(Fig.3B)**. The relative abundance of SARS-CoV-2 transcripts were similar in Calu-3 and infected hamster lung tissue. Analysing samples from the infected Calu-3 cells by northern blotting confirmed that FG-4592 treatment reduced viral transcripts **(Fig.3C)**. Furthermore, FG-4592 treatment inhibited N protein expression in infected Calu-3 cells **(Fig.3D)**, providing an explanation for the antiviral activity of HIFs.

**Figure 3:**
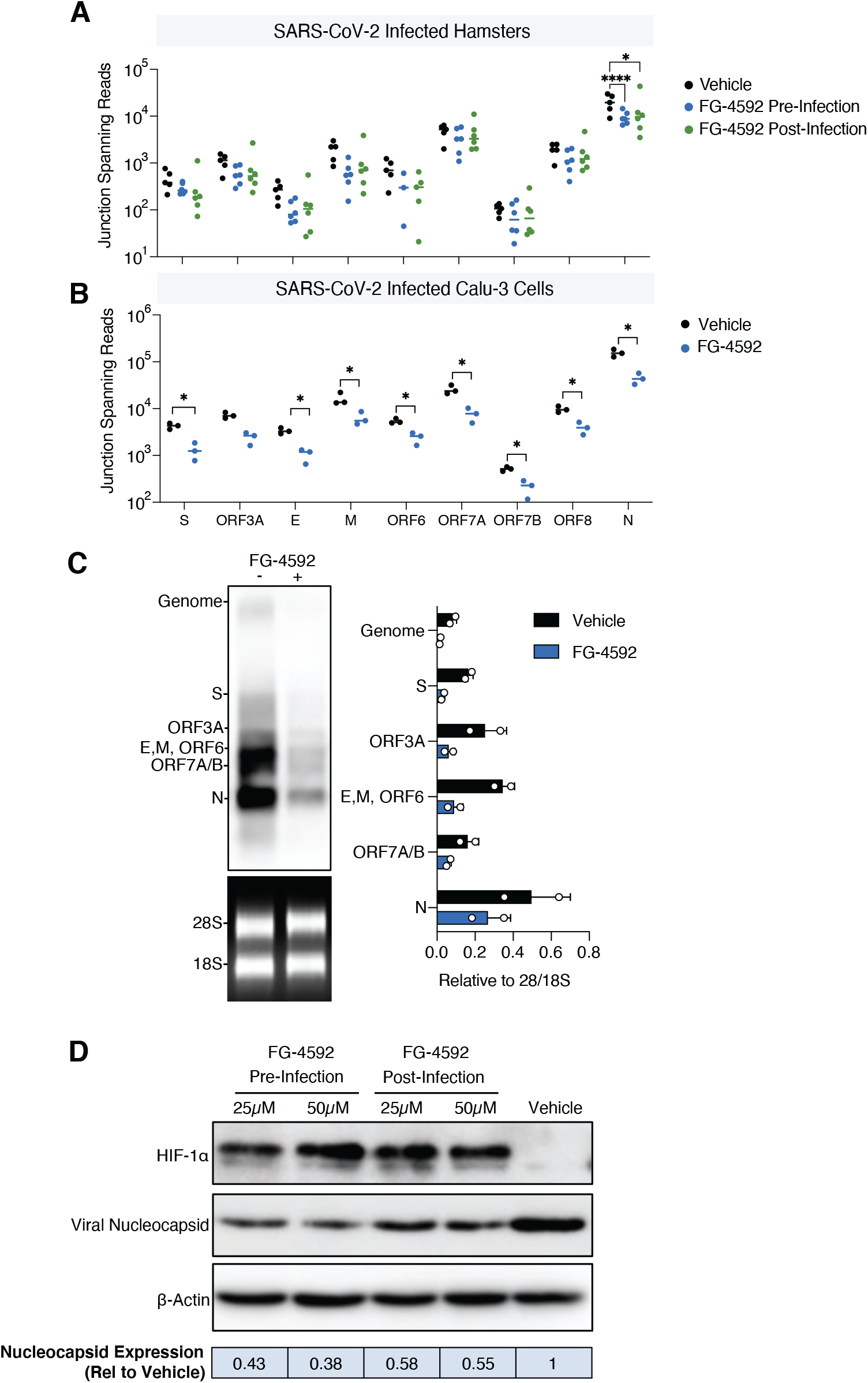
Analysis of the SARS-CoV-2 transcriptome in FG-4592 treated hamster lung and in Calu-3 cells. **(A)** Viral sequencing reads from infected lung tissue were mapped to the SARS-CoV-2 Victoria reference genome to generate a read depth profile. Reads that spanned the canonical viral sub-genomic transcript junctions were quantified for each tissue sample. **(B)** Junction spanning reads quantified from control or FG-4592 (50μM) pre-treated Calu-3 cells infected with SARS-CoV-2 (MOI of 0.01) (N=3 biological replicates). **(C)** Viral RNAs from the experiment in (B) were resolved on an RNA-agarose gel and analysed by Northern Blot hybridisation using a dioxygeninin labelled probe designed to detect all viral transcripts. Relative expression of transcripts was determined by densitometric analysis relative to the 28S/18S ribosomal RNA. **(D)** Representative immunoblot of SARS-CoV-2 nucleocapsid (N) and HIF-1α expression in Calu-3 cells treated with FG-4592 (25 or 50μM) pre- or post-infection. N expression was quantified by densitometry relative to β-actin and data expressed relative to the vehicle control. Statistical analysis was performed by ANOVA, p<0.05 = *, p<0.0001 = ****. Brackets indicate the comparisons tested.

Finally, we assessed SARS-CoV-2 sequence variation to determine whether treatment associated with genetic changes. Viral sequences were conserved across the genome in the vehicle or treated lung tissues **(Supplementary Fig.4B)** and no changes in the consensus sequence were seen in the treated animals or Calu-3 cells, with 100% conservation of the nucleotide sequence across the genome **(Supplementary Table 2)**. Together these data show that FG-4592 treatment had no effect on the sequence of SARS-CoV-2 in the lung but reduced sgRNAs.

### Spatial analysis of SARS-CoV-2 RNA and nucleocapsid expression in the respiratory tract

As FG-4592 reduced the clinical signs and levels of infectious virus in the upper and lower respiratory tract we explored the impact of treatment on virus-associated pathology. Sequential sections from the nasal cavity and lung tissue were stained with haematoxylin and eosin (H&E) and with RNA-scope *in situ* hybridization (ISH) probes targeting the Spike gene to assess the tissue distribution of SARS-CoV-2 RNA. We noted extensive inflammatory cell exudate in the nasal cavity and mild to moderate necrosis in both the olfactory and respiratory epithelia **(Supplementary Fig.5A-C)**. We assessed these pathological changes using a semi-quantitative scoring system and showed a reduction in the nasal histopathological score in the post-infection treatment group **(Fig.4B, Supplementary Table 3)**. Viral RNA primarily localised to the epithelia and exudate in the nasal cavity and FG-4592 reduced epithelial staining, the major site of virus replication **(Fig.4B)**. A similar observation was noted for viral RNA signals in the exudate **(Fig.4B);** however, these results may be compromised by the daily collection of nasal washes.

**Figure 4:**
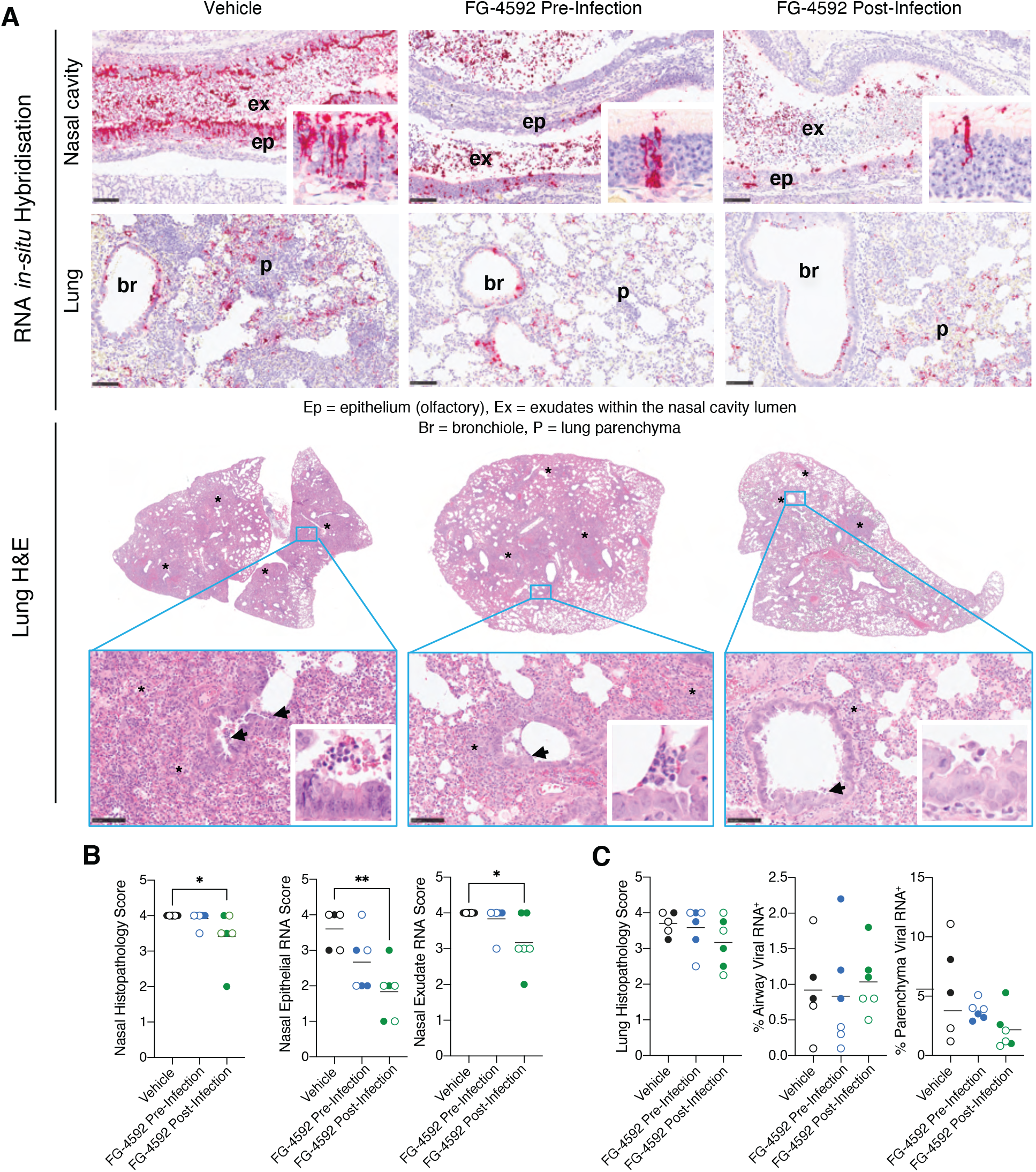
SARS-CoV-2 RNA and pathology in the nasal tract and lung. **(A)** Sections from the left lung and nasal cavity were stained with H&E to analyse histopathological changes and with RNA *in-situ* hybridisation (ISH) to detect SARS-CoV-2 RNA using probes specific for the viral Spike (S) transcript. Representative images from each treatment group are shown. Lung lesions consisted of multifocal broncho-interstitial pneumonia (*) with mild to moderate necrosis of the bronchiolar epithelium (arrows and H&E inserts) (see **Supplementary Fig.5**). ISH shows abundant viral RNA in the nasal cavity epithelia that associates with mild to moderate necrosis (inserts) and within the airway epithelia and areas of inflammation. Scale bar = 100μm. **(B)** Nasal cavity was assessed for the presence and severity of lesions using a semi-quantitative scoring system from H&E-stained sections. SARS-CoV-2 RNA in the olfactory epithelium or exudates was quantified using the following scoring system: 0=no positive staining; 1=minimal; 2=mild; 3=moderate and 4=abundant staining. **(C)** Lung pathology was assessed for the presence and severity of lesions using a semi-quantitative scoring system from H&E-stained sections. Digital image analysis calculated the area of lung airway and parenchyma staining for viral RNA. Statistical analysis was performed using a one-way ANOVA, p<0.05 = *, p<0.01 = **. N=5 animals for vehicle and N=6 animals for each treatment group.

In agreement with previous studies (Dowall et al., 2021; Gruber et al., 2020; Nouailles et al., 2021), SARS-CoV-2 infected lung tissue showed pulmonary lesions consisting of broncho-interstitial pneumonia extending into the alveoli and multifocal areas of consolidation, consistent with inflammatory cell infiltration and oedema **(Fig.4A, Supplementary Fig.5)**. Digital image analysis showed that FG-4592 treatment did not alter the severity of lung histopathology **(Fig.4C)**. Viral RNA was detected in the bronchiolar epithelia, bronchiolar inflammatory exudates, as well as in the lung parenchyma of the control vehicle animals **(Fig.4A)** and FG-4592 treatment had no detectable effect on the viral RNA signals in the parenchyma or airways **(Fig.4C)**.

To extend these observations we stained the infected nasal and lung sections for SARS-CoV-2 N antigen expression by immunohistochemistry (IHC). Within the nasal cavity, N primarily localised to the epithelia and exudate **(Fig.5A)**, consistent with the detection of S-gene transcripts. Semi-quantitative scoring of the nasal cavity sections showed reduced N antigen expression in the treated animals, most notably in the epithelia of post-infection treated samples **(Fig.5B)**. Within the lung, N staining localised to the airways and lung parenchyma **(Fig.5C)**, and we noted a significant reduction of parenchymal staining in the post-infection treated animals **(Fig.5D)**. In summary, histopathological analysis shows that PHI treatment reduced SARS-CoV-2 RNA and N antigen levels in the nasal epithelia and exudate, consistent with the reduction in infectious viral burden.

**Figure 5:**
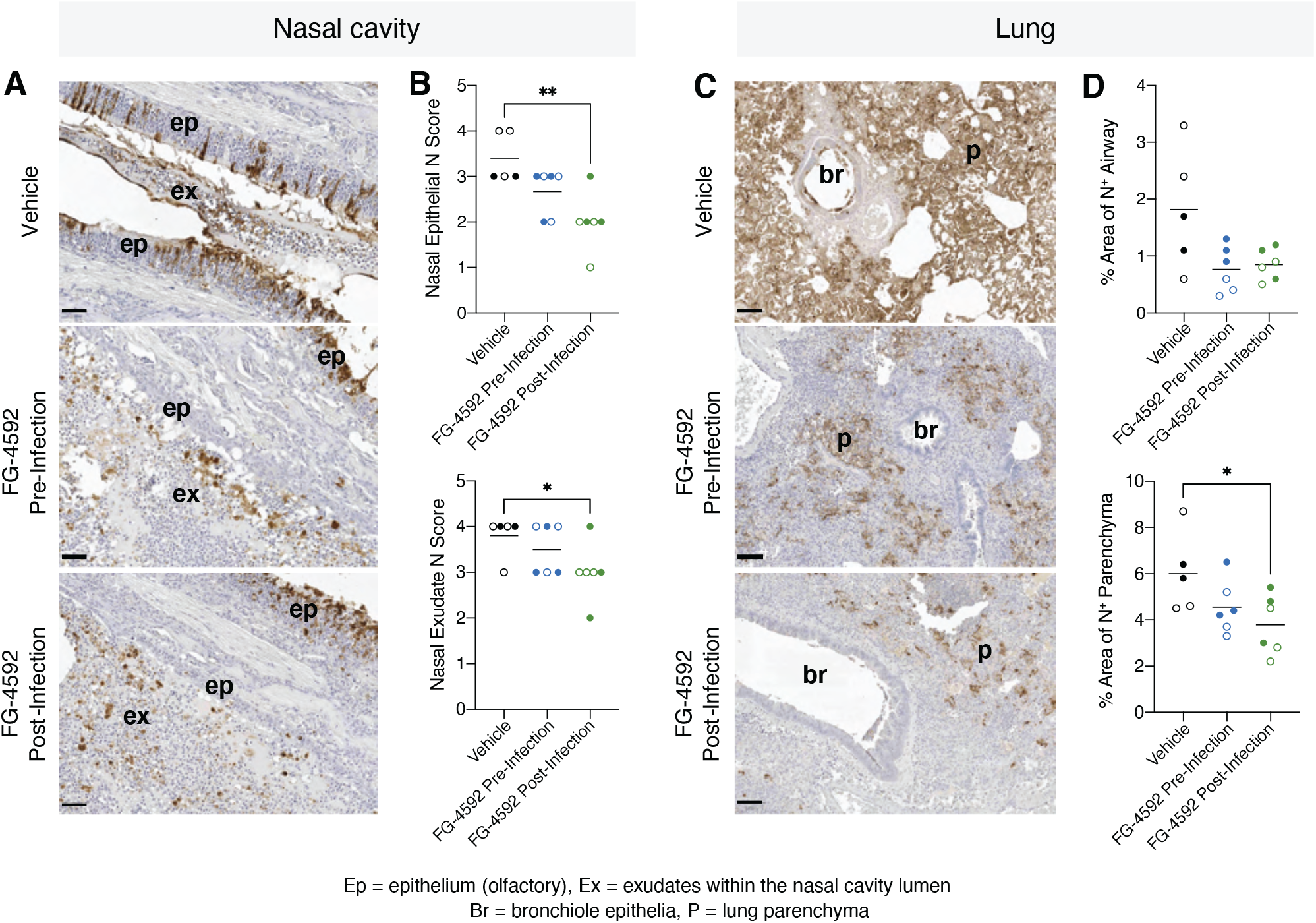
SARS-CoV-2 nucleocapsid expression in the nasal tract and lung. **(A)** Sections from the nasal cavity were stained by immunohistochemistry for the presence of the SARS-CoV-2 N protein. Representative images from each treatment group are displayed. N protein staining was observed in the olfactory epithelia (ep) and the inflammatory exudates (ex) within the nasal cavity luminae. Bar = 50μm. **(B)** A semiquantitative scoring system evaluated N protein expression in the olfactory epithelia or exudates, where: 0=no positive staining; 1=minimal; 2=mild; 3=moderate and 4=abundant staining. **(C)** Lung sections were assessed for N protein expression and representative images show positive staining in the bronchiolar epithelia, luminae (br) together and parenchyma (p). **(D)** Digital image analysis calculated the area of lung airway and parenchyma staining for N protein. Statistical analysis was performed by ANOVA, p<0.05 = *, p<0.01 = **. N=5 animals for vehicle and N=6 animals for each treatment group.

### FG-4592 reduces ciliated epithelial damage in the lung

To understand the global host response, we sequenced lung tissue from SARS-CoV-2 infected (vehicle control) and uninfected hamsters. Infection induced substantial changes in the lung transcriptome compared to uninfected tissue; with an up-regulation of pathways involved in inflammation and down-regulation of genes involved in cilium organisation and assembly **(Fig.6A-B)**. Analysing inflammatory gene expression using genes from the molecular signature database (Liberzon et al., 2015) showed that FG-4592 had a modest effect on the lung inflammasome **(Supplementary Fig.6)**. Of note, expression of the viral entry receptors *Ace2* and *Tmprss2* were significantly downregulated (log_2_ fold change of −1.45 and −1.52 respectively) in the infected tissue, likely reflecting viral cytopathology (Cross et al., 2021). Loss of ciliated epithelial cells is a key feature of COVID-19 resulting from damage to the airway epithelia (Pizzorno et al., 2020; Robinot et al., 2021; Zhu et al., 2020). To examine whether FG-4592 treatment reduced the level of epithelial damage we used the reported compendium of cilia-related genes (van Dam et al., 2019) to evaluate gene expression in the pulmonary transcriptome of vehicle and FG-4592 treated animals. A similar pattern of cilia-related gene expression was noted in the vehicle and pre-infection treatment group, with most genes showing a marked down-regulation **(Fig.6C**). However, the virus induced down-regulation of ciliated gene expression was less apparent in animals treated with FG-4592 post-infection **(Fig.6D)**. To gain further insight as to whether FG-4592 treatment affects the level of ciliated cells we stained nasal and lung sections for α-tubulin and club cell secretory protein (CCSP), markers of ciliated cells and secretory cells, respectively. We observed a substantial reduction in α-tubulin and CCSP staining in the infected lung **(Fig.6E)** and nasal cavity **(Supplementary Fig.6)**, consistent with virus-induced loss of ciliated cells **(Fig.6E)**. While limited staining was observed in the lung sections from the pre-treated animals, we noted a restoration of α-tubulin expression in post-infection treated sections, in line with our transcriptomic analysis. Together these data show that a loss of ciliated cells in the respiratory tract is a prominent feature of SARS-CoV-2 infection and FG-4592 treatment may offer some protection from this severe pathological change.

**Figure 6:**
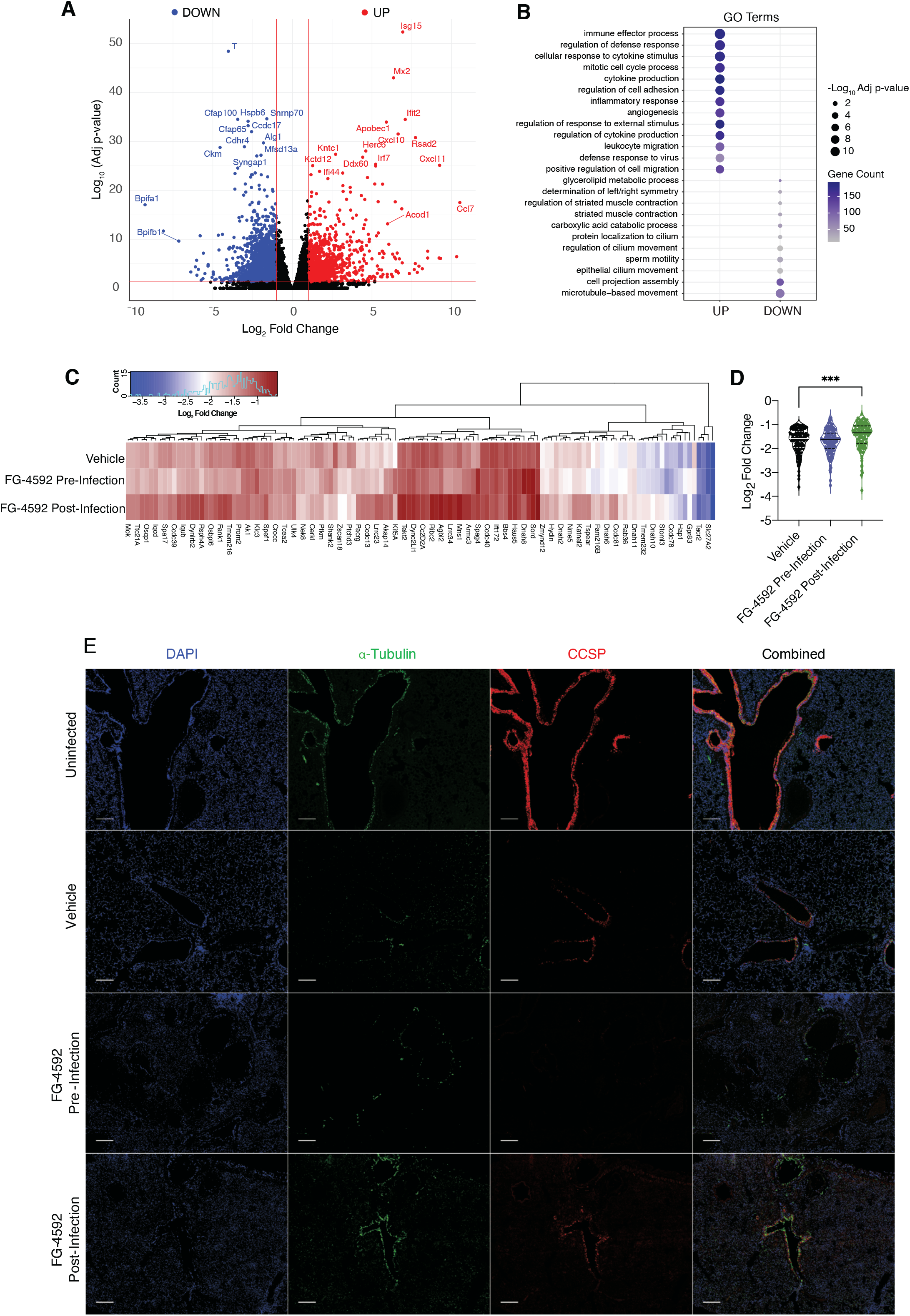
SARS-CoV-2 driven changes in lung ciliated epithelia. (**A**) RNA extracted from snap frozen lung tissue from, vehicle (N=5), FG-4592 treated pre- (N=6) and post-infection (N=6), or uninfected animals (N=2) was sequenced. Differential gene expression, defined as a log_2_ fold change of +/- 1 with an adjusted p value <0.05, was measured for the SARS-CoV-2 infected vehicle and uninfected samples. (**B**) Significant gene ontology pathways are displayed for both up and down regulated genes in infected vehicle vs uninfected samples, where symbol size reflects the -log10 adjusted p value and gene count is represented by the colour. (**C**) Expression of differentially expressed cilia-related genes, in vehicle, pre- and post-treatment groups compared to uninfected samples. Expression data is grouped by hierarchical clustering. (**D**) Log_2_ fold change values for all genes represented in C and grouped by treatment. Statistical analysis was performed using a one-way ANOVA, *** = p<0.001. (**E**) Representative lung sections stained for **α**-tubulin or CCSP from uninfected, vehicle, FG-4592 pre-or post-infection treated animals where nuclei are visualised with DAPI. Individual stains are shown along with an overlayed image. Scale bars represent 100μm.

## DISCUSSION

In this study we evaluated the antiviral potential of FG-4592 in the Syrian hamster model of SARS-CoV-2 infection. Treating animals pre- or post-infection reduced the levels of infectious virus and improved clinical symptoms. The drug was well tolerated, with no adverse reactions reported in any of the treated animals. Despite treatment showing a significant reduction in the levels of infectious virus, bulk PCR quantification of viral RNA in the nasal washes and throat swabs were unchanged. An earlier study of SARS-CoV-2 infection in Syrian hamsters reported a relatively short contagious period that associated with the detection of infectious virus (Dowall et al., 2021). However, SARS-CoV-2 RNA can persist in the respiratory tract long after the communicable period has passed (Chan et al., 2020; Dowall et al., 2021; Sia et al., 2020). SARS-CoV-2 RNA genomes are highly structured, and this may contribute to their persistence (Huston et al., 2021; Simmonds et al., 2021). Several reviews have reported a discrepancy between viral RNA levels and the detection of infectious virus in clinical samples (Cevik et al., 2021; La Scola et al., 2020; Walsh et al., 2020). While quantitative PCR measurement is the gold standard for SARS-CoV-2 diagnosis, this method only detects the viral nucleic acid and not the infectious capacity of virus particles. An important factor to consider is the cellular location of the viral RNA in the respiratory tract, where ISH probing of infected nasal tissue revealed SARS-CoV-2 RNA in both the nasal exudate and epithelia. We hypothesise the exudate will comprise extracellular encapsidated viral RNA and processed viral particles in immune cells and does not reflect sites of active virus replication. Our *in-situ* analysis highlights a significant reduction of viral RNA in the nasal epithelia of FG-4592 treated animals, demonstrating antiviral drug activity at the primary site of replication.

While the disconnect between viral RNA levels and infectivity is not well understood in a clinical setting, we would predict that drugs inhibiting the level of infectious virus in the upper respiratory tract will reduce virus transmission. The recently approved anti-viral drug Molnupiravir showed a negligible effect on viral RNA or infectious titre in respiratory samples when tested in Syrian hamsters (Rosenke et al., 2021). Yet both Molnupiravir and FG-4592 treatments were associated with significant reductions in the burden of infectious virus in the lung. While both drugs have contrasting mechanisms of action, the ability of FG-4592 to limit viral replication highlights the value of targeting host pathways that are essential for viral replication in concert with the development of direct-acting antiviral (DAA) agents. A promising area for future development is a combined treatment of PHI and DAAs such as Molnupiravir or the recently approved protease inhibitor Nirmatrelvir (Owen et al., 2021). Previous strategies of DAA monotherapies for the treatment of HIV and HCV selected for drug resistant viruses, reinforcing the value of combination therapies that result in a high barrier to the development of anti-viral resistance (Hiscox et al., 2021). Our analysis of the SARS-CoV-2 sequences showed no evidence of mutational change in treated animals. However, the viral transcriptome in the lung showed differences in the abundance of sgRNAs in the treated animals, similar to our observations with infected Calu-3 cells where FG-4592 significantly reduced sgRNAs and N protein expression. These data show a role for HIFs in regulating SARS-CoV-2 sgRNA levels that could be explained by changes in the genesis or maintenance of viral replication complexes, in line with our previous observations (Wing et al., 2021a).

An important finding of this study was the improved clinical score in the treated animals. FG-4592 substantially reduced the incidence of laboured breathing in the infected animals, irrespective of treatment grouping, that may be attributed to the increased levels of erythropoiesis resulting in improved blood oxygenation in the infected hamsters. These results justify future studies to evaluate how improving blood oxygenation impacts the clinical outcome of SARS-CoV-2 infection.

Histopathological analysis of pulmonary tissue showed that a short duration of FG-4592 treatment improved virus-induced pathology in the upper respiratory tract. Lungs from the treated animals showed fewer areas of consolidation and pulmonary lesions, particularly in the post-infection treatment group. An earlier study reported that SARS-CoV-2 induced lung pathology in this experimental model peaked at 5 days post-infection, after which tissue injury recovered along with viral clearance (Rosenke et al., 2020). Since the primary goal of this study was to evaluate the effect of PHIs on viral replication rather than pathological changes, we predict that longer PHI treatment times would have more pronounced effects on pulmonary pathology and potentially improve recovery times.

A key feature of SARS-CoV-2 infection is the profound loss of the ciliary layer in the respiratory tract (Pizzorno et al., 2020; Robinot et al., 2021; Zhu et al., 2020); induced either by direct infection of these cells and their subsequent dedifferentiation (Robinot et al., 2021) or loss by cytopathic infection. The resulting impairment in mucociliary clearance limits the removal of infiltrating viral, bacterial, or fungal pathogens that can lead to secondary infections with antibiotic resistant strains of *Staphylococcus aureus* and *Klebsiella pneumoniae* (Manohar et al., 2020), the incidence of which markedly increases in critically ill patients (Alanio et al., 2020; Manohar et al., 2020). Our lung transcriptomic analysis highlighted the altered expression of genes involved in ciliated cell function in FG-4592 treated animals. Components of the centrosome, important for cilia formation, are regulated by PHDs and hypoxia increases their expression (Moser et al., 2013), providing a potential mechanism for our observation. Nonetheless, we are unable to assess whether treatment prevents infection of ciliated cells or promotes their recovery. Staining tissue sections for α-tubulin expression, a marker for ciliated cells, in the upper and lower respiratory tract shows the considerable impact of infection on the abundance of ciliated cells. Importantly, lung sections from animals treated with FG-4592 post-infection showed elevated levels of ciliated cell staining, that may reflect differences in the transcriptional response of infected cells to PHIs. Healthy cells will trigger reactions enabling them to adapt to hypoxia such as switching from oxidative phosphorylation to glycolysis (Semenza, 2011), however, SARS-CoV-2 infection may alter the cellular response to hypoxia potentially exacerbating necrosis, cytokine expression and inflammatory responses (Serebrovska et al., 2020).

At present there are limited therapeutic options for treating COVID-19 most of which treat the clinical manifestation of the disease. FG-4592 treatment before or after SARS-CoV-2 infection reduced the infectious viral burden and restored the loss of ciliated cells, providing an opportunity to improve clinical outcomes by limiting secondary infections. Our observations may be applicable for the treatment of other respiratory pathogens, including both Influenza A virus (Ren et al., 2019; Zhao et al., 2020) and Respiratory Syncytial virus, whose replication has been reported to be HIF-dependent (Morris et al., 2020). In summary, we demonstrate a role for HIFs to suppress SARS-CoV-2 infection and associated disease in an experimental animal model, highlighting the value of prolyl-hydroxylase inhibitors for the treatment of COVID-19.

## METHODS

### Animals

Golden Syrian hamsters 9 (*Mesocricetus auratus*) aged 6–8 weeks were obtained from Envigo RMS UK Ltd., Bicester, UK. The animals were housed in cages that are designed in accordance with the requirements of the UK Home Office Code of Practice for the Housing and Care of Animal Used for Scientific Procedures (1986). During procedures with SARS-CoV-2, the animals were housed in a flexible-film isolator within a Containment Level 3 facility. The animals were randomly assigned into groups and individually housed, with equal allocation of male and female animals to each study. For direct intranasal challenge studies, group sizes of 6 hamsters were used as the minimal number required for statistical significance to be achieved. Access to food and water was ad libitum and environment enrichment was provided. Rooms were maintained within set parameters: 20–24 °C, 45–65% humidity and a 12/12 light cycle.

### Syrian Hamster study design and ethics approval

Animals were divided into three groups for treatment with vehicle, FG-4592 pre-infection or post-infection (n=6 per group). Animals were treated with 30mg/kg of FG-4592 (MedChem Express) by oral gavage. Drug was dissolved in 99% double distilled H_2_O, 0.5% methyl cellulase and 0.5% Tween-80, administered twice daily. Treatment commenced either 24h prior to (pre) or 24h following (post) infection and maintained until termination of the study (4 days) after infection. The control group followed the dosing schedule of the pre-infection group and were treated with vehicle only. All groups were infected intranasally with 5×10^4^ PFU of Australia/VIC01/2020 SARS-CoV-2. Viral inocula were made in sterile phosphate buffered saline (PBS) and delivered via intranasal instillation (200μL total with 100μL per nare) with animals sedated using isoflurane. Animal weights and temperatures were monitored daily and visual inspection of all animals carried out twice daily, with signs of clinical disease such as wasp waisted, ruffled fur, hunched or laboured breathing recorded (**Supplementary Table 1**). Throat swabs and nasal washes were collected on days 1, 2 and 4 post infection. Animals were euthanised at day 4 post infection and tissues collected at necropsy for pathology and virology assays. All experimental work was conducted under the authority of a UK Home Office approved project licence that had been subject to local ethical review at Public Health England (now part of the UK Health Security Agency (UKHSA) Porton Down by the Animal Welfare and Ethical Review Body (AWERB) as required by the Home Office Animals (Scientific Procedures) Act 1986.

### Virus and cells

SARS-CoV-2 Australia/VIC01/2020(Caly et al., 2020) was provided by the Peter Doherty Institute for Infection and Immunity, Melbourne, Australia at P1 and passaged twice in Vero/hSLAM cells (Cat#04091501) obtained from the European Collection of Cell Cultures (ECACC), UK. Virus infectivity was determined by plaque assay on Vero-TMPRSS2 cells as previously reported (Wing et al., 2021a). Calu-3 cells were obtained from Prof Nicole Zitzmann’s lab and maintained in Advanced DMEM, 10% FCS, L-glutamine and penicillin streptomycin. Calu-3 cells were infected with the above strain of SARS-CoV-2 at an MOI of 0.01 for 2h. Viral inocula were removed, cells washed three times in PBS and maintained in growth media until harvest.

### Plaque assay quantification of virus infectivity

Samples from nasal washes, throat swabs or lung homogenates were serially diluted 1:10 and used to inoculate monolayers of Vero-TMPRSS2 cells for 2h. Inocula were removed and replaced with DMEM containing 1% FCS and a semi-solid overlay consisting of 1.5% carboxymethyl cellulose (SIGMA). Cells were incubated for 72h, after which cells were fixed in 4% PFA, stained with 0.2% crystal violet (w/v) and visible plaques enumerated.

### qPCR quantification

Viral RNA was extracted from nasal washes or throat swabs using the QiaAMP Viral RNA kit (Qiagen) according to manufacturer’s instructions. Tissues were homogenised using the GentleMACS homogeniser in RLT buffer and extracted using the RNeasy kit (Qiagen) according to manufacturer’s instructions. For quantification of viral or cellular RNA, equal amounts of RNA, as determined by nanodrop, were used in a one-step RT-qPCR using the Takyon-One Step RT probe mastermix (Eurogentec) and run on a Roche Light Cycler 96. For quantification of viral copy numbers, qPCR runs contained serial dilutions of viral RNA standards. Total SARS-CoV-2 RNA was quantified using: 2019-nCoV_N1-F: 5’-GAC CCC AAA ATC AGC GAA AT-3’, 2019-nCoV_N1-R: 5’-TCT GGT TAC TGC CAG TTG AA TCT G-3’, 2019-nCoV_N1-Probe: 5’-FAM-ACC CCG CAT TAC GTT TGG TGG ACC-BHQ1-3’. Genomic viral RNA was quantified using SARS-CoV-2-gRNA_F: 5’-ACC AAC CAA CTT TCG ATC TCT TGT-3’, SARS-CoV-2-gRNA_R: 5’-CCT CCA CGG AGT CTC CAA AG-3’, SARS-CoV-2-gRNA_Probe: 5’ FAM-GCT GGT AGT GAC TGC TTT TCG CCC C-BHQ1-3’. Hamster host transcripts were quantified using the following Taqman expression assays by ThermoFisher, *Edn1* (APAAFZZ), *Ace2* (1956514) and *β-Actin* (APZTJRT).

### Histopathology, *in situ* hybridisation and Immunohistochemistry

The nasal cavity and left lung were fixed by immersion in 10% neutral-buffered formalin and processed into paraffin wax. Nasal cavity samples were decalcified using an EDTA-based solution prior to longitudinal sectioning to expose the respiratory and olfactory epithelium. Sequential 4 μm sections were stained with H&E. In addition, samples were stained using the *in-situ* hybridisation (ISH) RNAscope technique to label SARS-CoV-2 RNA using V-nCoV2019-S probe (Cat No. 848561, Advanced Cell Diagnostics). Briefly, tissues were pre-treated with hydrogen peroxide for 10 min (room temperature), target retrieval for 15 min (98–101°C) and protease plus for 30 min (40°C) (Advanced Cell Diagnostics). The probe was incubated with the tissues for 2h at 40°C and the signal amplified using RNAscope 2.5 HD Detection kit – Red (Advanced Cell Diagnostics). Immunohistochemical (IHC) staining of the SARS-CoV-2 nucleocapsid (N) protein, deparaffinisation and heat-induced epitope retrieval were performed on the Leica BOND-RXm using BOND Epitope Retrieval Solution 2 (ER2, pH 9.0) for 30 minutes at 95°C. Staining was performed with the BOND Polymer Refine Detection kit, a rabbit anti-SARS-CoV-2 nucleocapsid antibody (Sinobiological; clone: #001; dilution: 1:5000) and counterstained with haematoxylin. The H&E, ISH and IHC stained slides were scanned using a Hamamatsu S360 digital slide scanner and examined using ndp.view2 software (v2.8.24). Lung tissue from one animal in the vehicle group was not processed due to deterioration of the sample. Digital image analysis using Nikon NIS-Ar software quantified SARS-CoV-2 RNA or N expression in the lung sections by calculating the percentage of positively stained areas in defined regions of interest (ROI), including the airway epithelia and parenchyma. For the nasal cavity a semi-quantitative scoring system (Dowall et al., 2021) evaluated the presence of SARS-CoV-2 RNA or N expression in the exudate and epithelia where: 0=no staining; 1=minimal; 2=mild; 3=moderate and 4=abundant staining. All slides were evaluated subjectively by a qualified pathologist, blinded to treatment details and were randomised prior to examination to limit bias (blind evaluation). Random slides were peer-reviewed by a second pathologist. Histopathology was carried out in a ISO9001:2015 and GLP compliant laboratory. A semiquantitative scoring system evaluated the severity of lesions in the lung and nasal cavity as previously reported (Dowall et al., 2021).

### RNA sequencing and data analysis

RNA was extracted from 30mg of homogenised lung (right lobe) using the RNeasy kit (Qiagen) and RNA integrity determined by Tapestation (Agilent), before providing RNA to Novogene UK Ltd for poly-A-enriched transcriptome sequencing. Paired end Illumina sequencing was carried out with a300bp fragment length and mapped to the *Mesocricetus auratus* genome. Viral reads were mapped to the SARS-CoV-2 reference genome (NC 045512.2) using Salmon (Patro et al., 2017). FPKM values were enumerated, and differential expression quantified using the DeSeq2 package (Love et al., 2014). Threshold for statistical significance was set as log_2_FC +/- 1 and an adjusted p value <0.05. Junction spanning reads were detected as described in Kim et al 2021(Kim et al., 2020) using the ggsashimi analysis package (Love et al., 2014).

### Immunoblotting

Cells were prepared by washing cells with PBS and lysed using RIPA buffer (20 mM Tris, pH 7.5, 2 mM EDTA, 150 mM NaCl, 1% NP40, and 1% sodium deoxycholate) supplemented with protease inhibitor cocktail tablets (Roche). Clarified samples were mixed with laemmli sample buffer, separated by SDS-PAGE and proteins transferred to polyvinylidene difluoride membrane. Membranes were blocked in 5% milk in PBS/0.1% Tween-20 and incubated with anti-HIF-1α (BD Biosciences), anti-β-Actin (Sigma) or SARS-CoV-2 nucleocapsid (EY-2A, a kind gift from Prof Alain Townsend) primary antibodies and appropriate HRP-conjugated secondary antibodies (DAKO). Chemiluminescence substrate (West Dura, 34076, Thermo Fisher Scientific) was used to visualize proteins using a G:Box Imaging system (Syngene).

### Visualising SARS-CoV-2 RNAs by Northern Blotting

Infected Calu-3 cells were harvested in Trizol (Thermofisher) 24h post infection and total RNA extracted according to manufacturer’s instructions. 10 μg of RNA was resolved on a 10% MOPS, 2.2 M formaldehyde agarose gel. To show equal RNA loading, the 18S and 28S ribosomal subunit RNA species were visualised under UV light through ethidium bromide staining. Gels were denatured in 50 mM NaOH for 5 minutes, and RNAs transferred to nylon membrane by capillary transfer in 1XSSC buffer. Membranes were washed and RNAs fixed by UV crosslinking. Membranes were hybridised at 65°C overnight with a digoxigenin-labelled DNA probe specific to the 3’ end of the SARS-CoV-2 genome, enabling the detection of all viral RNAs. Bands were visualised using a luminescent DIG detection kit (Roche) according to manufacturer’s instructions.

### Statistical Analysis

All data are presented as mean values ± SEM. P values were determined using the Mann-Whitney test (two group comparisons) or with the Kruskal–Wallis ANOVA (multi group comparisons) using PRISM version 8. In the figures * denotes p < 0.05, ** < 0.01, *** <0.001 and **** <0.0001.

## Data availability

The authors declare that all data supporting the findings of this study are available within the article and its Supplementary Information files or are available from the authors upon request. RNAseq data from this study are deposited on NCBI under the GEO accession ID: GSE195879 (https://www.ncbi.nlm.nih.gov/geo/query/acc.cgi?acc=GSE195879)

## ACKNOWLEDGEMENTS

The authors would like to thank our colleagues at the University of Oxford, Anderson Ryan and Nicole Zitzmann for Calu-3 cells, William James for Vero-E6 TMPRSS2 and Alain Townsend for anti-nucleocapsid. We acknowledge the support from the Biological Investigations Group and Histopathology Department at the UK Health Security Agency, Porton Down. JAM is funded by a Wellcome Investigator Award 200838/Z/16/Z, UK Medical Research Council (MRC) project grant MR/R022011/1 and Chinese Academy of Medical Sciences (CAMS) Innovation Fund for Medical Science (CIFMS), China (grant number: 2018-I2M-2-002). FI is funded by the Wellcome Trust 211122/Z/18 and AC is supported by an Oxford-BMS Fellowship. TB is funded by the Paradifference Foundation and COVID-19 Research Response Fund, University of Oxford. SC and AW are funded by the Francis Crick Institute, which receives its core funding from Cancer Research UK (FC001206), the UK Medical Research Council (FC001206), and the Wellcome Trust (FC001206).

## AUTHOR CONTRIBUTIONS

PACW designed and conducted experiments and co-wrote MS; MPB designed and conducted experiments; AC designed and conducted experiments; SC designed and conducted experiments; COR provided technical help; XC conducted experiments; JMH analysed data; XZ conducted experiments; RLJ performed experiments; KAR performed experiments; YH co-designed the study;; MWC helped with study design;; FI advised on tissue staining;; PB analysed data;; AW designed experiments and co-wrote MS; TB co-designed the study and co-wrote MS; FJS analysed data and co-wrote MS; JAM designed the study and co-wrote MS.

